# Diversity of vaccination-induced immune responses can prevent the spread of vaccine escape mutants

**DOI:** 10.1101/2021.04.14.439790

**Authors:** Judith A Bouman, Céline Capelli, Roland R Regoes

## Abstract

Pathogens that are resistant against drug treatment are widely observed. In contrast, pathogens that escape the immune response elicited upon vaccination are rare. Previous studies showed that the prophylactic character of vaccines, the multiplicity of epitopes to which the immune system responds within a host, and their diversity between hosts delay the evolution and emergence of escape mutants in a vaccinated population. By extending previous mathematical models, we find that, depending on the cost of the escape mutations, there even exist critical levels of immune response diversity that completely prevent vaccine escape. Furthermore, to quantify the potential for vaccine escape below these critical levels, we propose a concept of *escape depth* which measures the fraction of escape mutants that can spread in a vaccinated population. Determining this escape depth for a vaccine could help to predict its sustainability in the face of pathogen evolution.

## Introduction

Both drug treatment and vaccination induce an evolutionary pressure on pathogens. However, while drug-resistant pathogens are widely observed, pathogens that evolved the capability to escape the immune response primed by a vaccine are rare [1]–[4]. Vaccines, in contrast to drug treatment, work prophylactically, which reduces the probability that an escape mutant emerges by de novo mutation compared to drug treatment [1]. Additionally, vaccines work indirectly by eliciting immune responses, and thus profit from the characteristics of a fine-tuned system that has evolved over millions of years to prevent and clear infections [5], [6].

The immune responses elicited upon vaccination differ from person to person [7], whereas drug treatment often targets the same and a limited number of pathogen functions within each individual. The diversity of immune responses appears on two levels: the various epitopes to which the immune system responds within a vaccinated individual and the diversity of these epitopes between individuals. An epitope is the part of a antigen to which an antibody attaches, often, a single mutation in the epitope can confer escape from specific antibodies [8], [9]. The diversity in epitopes can be observed by studying antibody repertoires elicited by individuals upon vaccination, which recently became possible due to novel bioinformatical and statistical methods [10]. Such studies have shown that the antibody repertoires elicited after a tetanus vaccination, that exists of up to 100 clones [1], [11], barely overlap between individuals [1], [12]. Similarly, in a study of a pneumococcal polysacharide vaccination, all four donors elicited an antibody repertoire with an unique fingerprint [13]. Besides being based on factors such as age [14], [15], sex [16] and genetic background [17]–[19], twin studies have shown that this diversity is partly stochastic [20], [21].

The multiplicity of epitopes to which the immune system responds within a host increases the number of mutations needed to escape its immune system [1]. The between-host diversity in immune responses hinders escape mutants to spread in the population once they have emerged [1]. Kennedy and Read (2017) show, based on a mathematical model, that both, the prophylactic character and multiplicity of therapeutic targets delay the expected time at which escape mutants can emerge and have a synergistic effect when they are combined [1]. We aim to go beyond Kennedy and Read by specifying the conditions of the diversity in immune responses upon vaccination that can prevent escape mutants to spread at all. Moreover, we aim to understand the relative role of the multiplicity of immune responses against epitopes within an individual compared to their diversity between individuals.

To find conditions that prevent the spread of vaccine escape mutants, we use a mathematical model. This mathematical model includes a multiplicity of within-host immune responses against epitopes of the pathogen and between-host diversity of these responses (see Figure 1). Besides the wild-type pathogen, the model also includes escape mutants. These escape mutants contain one or multiple mutations in epitopes which allow them to escape the immune response an individual elicits against these epitopes. To capture the complex dynamics of the wild-type and multiple escape strains, we propose a new concept: the *escape depth*, which we define as the fraction of possible escape mutants that can spread in a vaccinated population. The escape depth allows us to quantify the ability of the pathogen to escape from a vaccine on a continuous scale and identify parameter values for which a vaccine is *evolution proof*.

**Figure 1:**
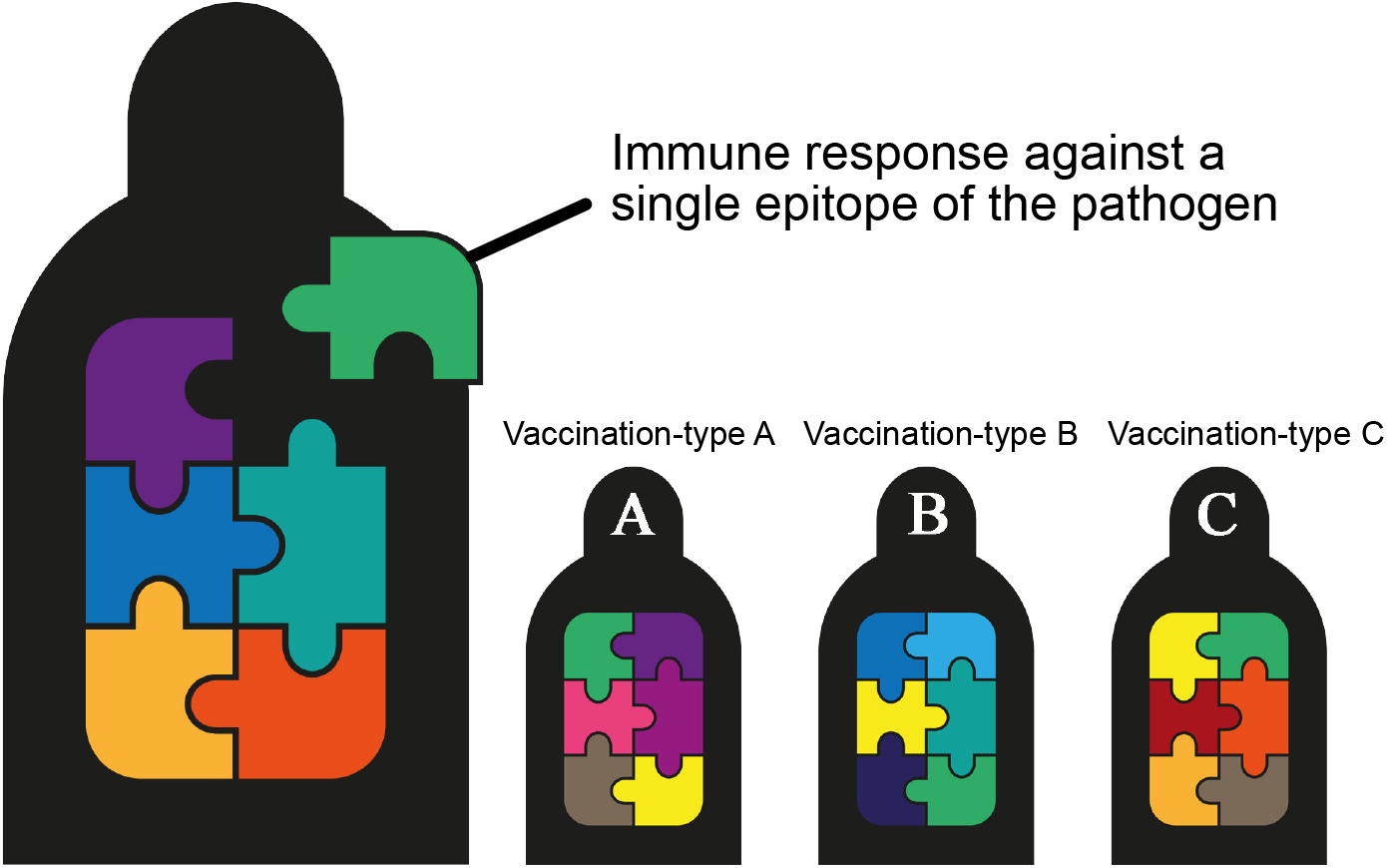
Representation of the modelled diversity of immune responses within and between individuals. Upon vaccination, each individual can elicit an immune response against epitopes of the pathogen selected from a larger set of epitopes. Every unique combination of immune responses is referred to as a vaccination-type. © M. Baldachin. Scientific illustrator. https://www.mishabaldachin.com.

## Results

To investigate the conditions under which the spread of an escape mutant in a vaccinated population is delayed or prevented, we extended McLean’s model (1995) to include within-host multiplicity and between-host diversity in the epitopes to which the immune system responds [22]. In our model, a vaccinated individual elicits an immune response against epitopes selected from a larger set that represents the total diversity in targeted epitopes. Each unique combination of responses against epitopes is referred to as a vaccination-type (see Figure 1). Pathogen mutants can escape the response against an epitope by a single mutation in this epitope. The cost of such an escape mutation is modelled by a decrease in the transmission rate of the mutant. To be precise, the transmission rate decreases with 10% for each additional mutation. Similar to the original McLean model, we assume that all possible escape mutants are present in the population at low levels before vaccination starts. Also, we assume that the vaccine has an efficacy of only 20% for preventing infection with an escape mutant.

Figure 2 shows the dynamics of the model for a single immune response per individual and either two or eight different responses between individuals (vaccination-types). This Figure illustrates what has been described by Kennedy and Read before: increasing the diversity in immune responses upon vaccination delays the expected rise of escape mutants. Here we investigate this qualitative pattern in more depth.

**Figure 2:**
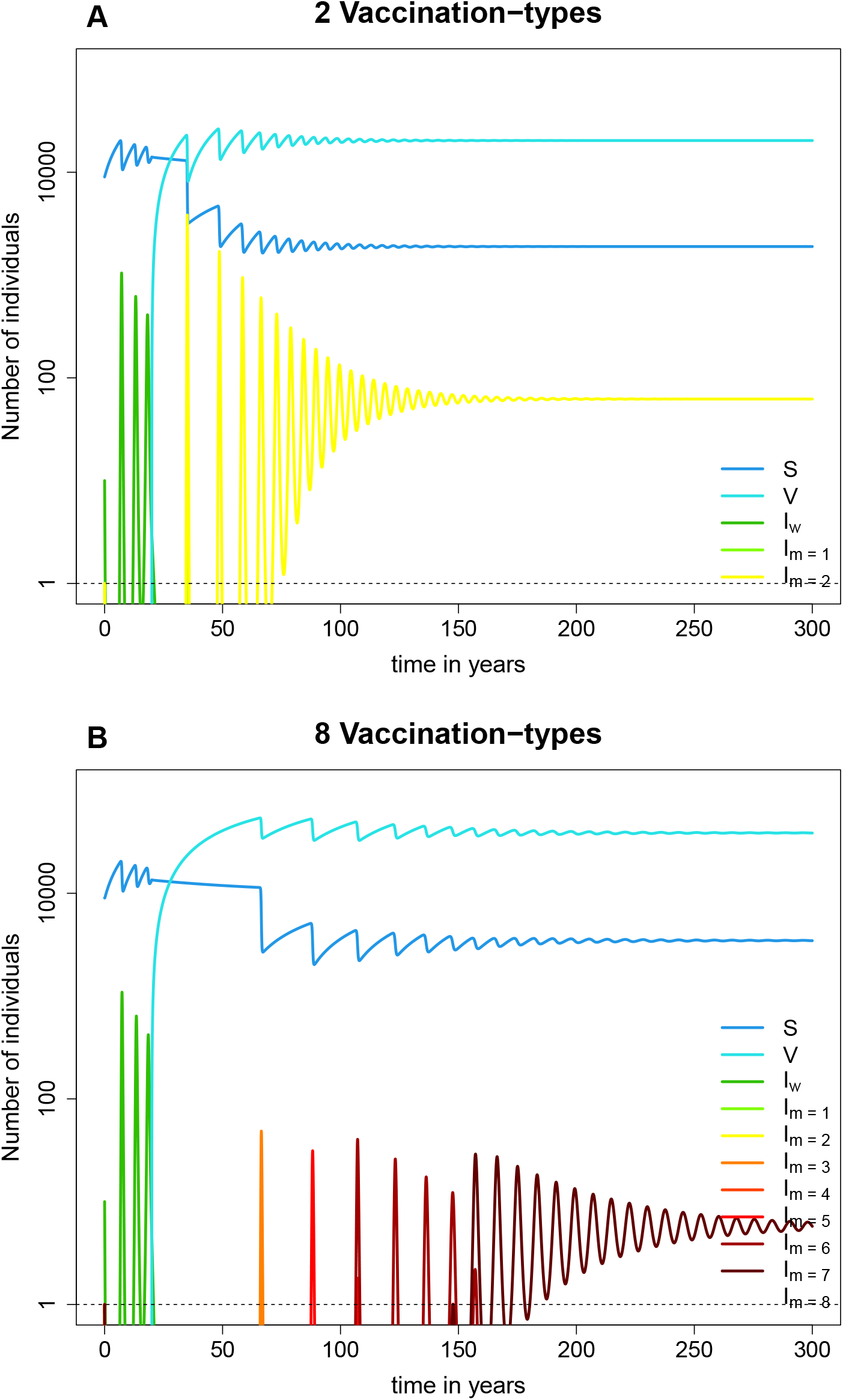
The dynamics of the mathematical model for a single within-host immune response and (A) 2 vaccination-types or (B) 8 vaccination types. Both panels show the number of non-vaccinated susceptibles (dark blue), vaccinated susceptibles (light blue), individuals infected with wild type pathogen (dark green) and individuals infected with one of the escape mutants. The number of individuals infected with an escape mutant are colored based on the number of mutations in the escape variant: light green for a single mutation to dark red for 8 mutations.

In Figure 2, we evaluated the model for a finite time. To determine whether an escape mutant could potentially spread at a later time we calculated the reproduction number for each mutant type. The reproduction number is the expected number of secondary cases due to one primary infected individual. A reproduction number larger than one will enable the mutant to spread in the population. The reproduction number of an escape mutant in a completely susceptible population without any other pathogens is referred to as the basic reproduction number (*R*_0_). The reproduction number of an escape mutant in a population where all other types of mutants with fewer escape mutations have already had the opportunity to spread is referred to as the endemic reproduction number (*R_e_*).

In general, the reproduction number can be calculated by taking the product of the transmission rate of the pathogen, the average infectious period and the number of susceptibles that can be infected. In our model, we assume *m* escape mutations resulting 2*^m^* escape mutant strains that each have their own reproduction number. Because escape mutations carry a transmission cost, the escape mutants differ in their transmission rate, *β_m_*. The average infectious period in our model is given by the inverse of the clearance rate *δ*, which we assume to be the same for each pathogen strain.

Lastly, the basic reproduction numbers of the pathogen strains are affected by the diversity of immune responses elicited by the vaccine within and between the hosts. To capture this diversity, we define a set of 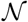 vaccination-types, and the escape matrix *J_n,m_* which equals one if mutant *m* can escape vaccination-type 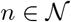 and is zero otherwise. We assume that escape mutations do not make a strain completely refractory the a given immune response. If a strain carries all mutations required to escape the immune responses within a vaccinated host it can infect it by a factor *r* less than an unvaccinated host. Without any escape mutations a vaccinated host is by a factor *s > r* less susceptible to infection.

The calculation of reproduction numbers relies on the levels of the number of unvaccinated and vacci-nated susceptibles, 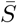 and 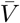, respectively. The basic reproduction number *R*_0_ can be calculated by using the levels of 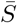 and 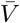 in the disease-free equilibrium, while the endemic reproduction number *R_e_* requires their levels endemic equilibrium.

### Between-host diversity of immune responses

Figure 4**A** and **B** show that the reproduction numbers of the escape mutants decrease for increasing between-host diversity of immune-responses with only one response within individuals. The basic reproduction number is shown for up to 70 vaccination types, where the endemic reproduction is limited to 11 vaccination types due to computational limitations. To simplify the interpretation of the many reproduction numbers shown in Figure 4**A** and **B**, we introduced the concept of *escape depth*. The escape depth is defined as the fraction of possible escape mutants that has a reproduction number above 1 and could thus spread in the vaccinated population (see Figure 4**C** and **D**). With an escape depth of 1 each possible mutant could spread, whereas an escape depth of 0 indicates that none of the mutants could spread. This last case would imply that the vaccine is completely evolution-proof (in our model only). The continuous scale between these two extremes helps to quantify and compare the evolutionary sustainability of vaccines. Figure 4**C** and **D** show that the escape depth of the model decreases with an increase of the number of vaccination types. Starting from 37 vaccination types, the escape depth based on the basic reproduction number is even zero, meaning that the vaccine can not be escaped for these levels of between-host diversity.

**Figure 3:**
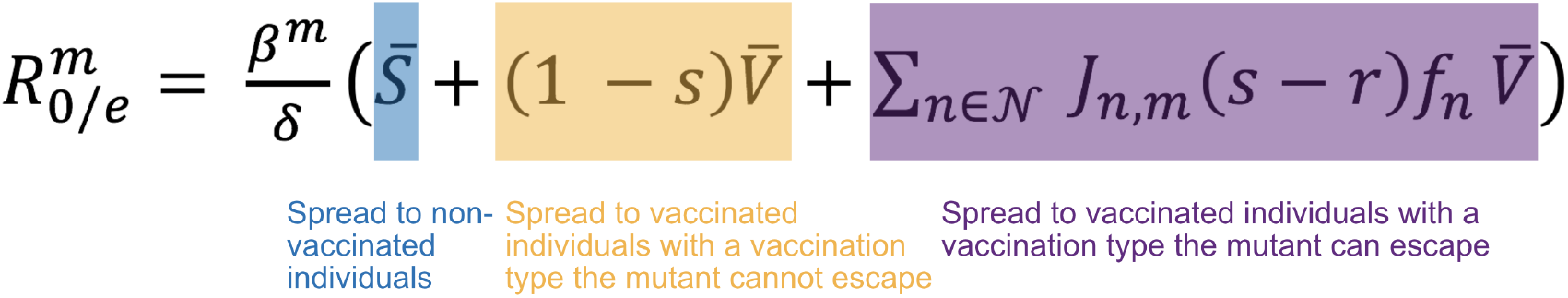
Equation for the reproduction number (either basic (*R*_0_) or endemic (*R_e_*)) of an escape mutant with *m* mutations. The constant 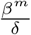 is the product of the transmission rate and the infectious time. This is multiplied with the number of individuals that can get infected by the pathogen type: unvaccinated susceptibles (blue), vaccinated individuals which can get infected by all pathogens with an efficacy of (1 − *s*) (yellow), and the vaccinated individuals with an immune response that this pathogen can escape and infect with efficacy (*s* − *r*).

**Figure 4:**
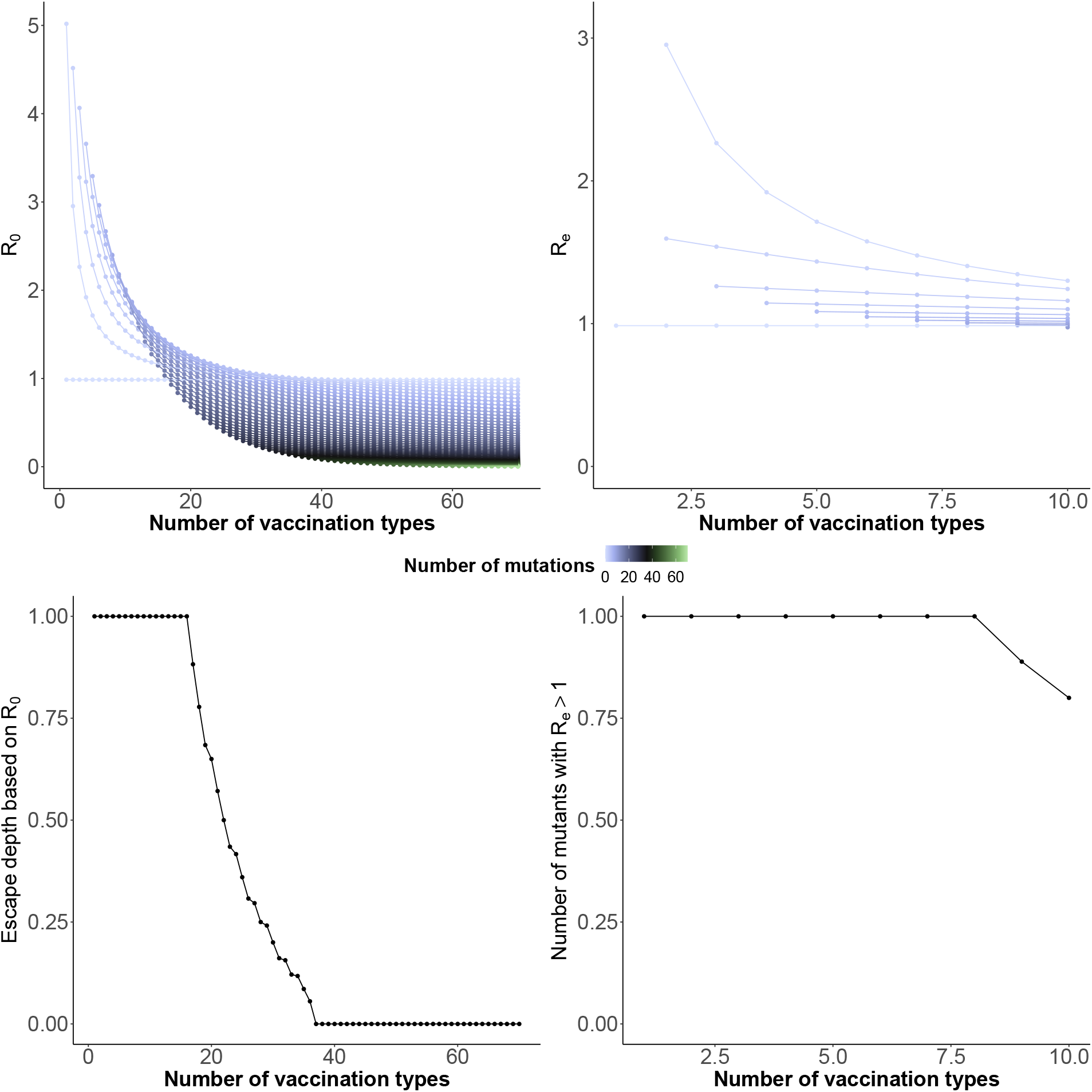
(A) The basic reproduction number and (B) the endemic reproduction number for escape mutants with varying number of escape mutations (ranging from 0 (light violet) to 70 (light green)) for increasing between-host immune diversity while there is a single immune responses within each individual. (C) The escape depth, i.e. the fraction of the escape mutants that has the potential to spread, against the number of vaccination types in the model. (D) Escape depth of the model based on the endemic reproduction number against the number of vaccination types in the model.

### Effect of model parameters on the escape-depth

The escape depth based on the basic reproduction number allows us to analyze the sensitivity of the model to three important model parameters. First, we evaluate the effect of the efficacy of the vaccine against the escape mutants. In the standard model, this value is fixed to 20%. In Figure 5**A** the value ranges from 0% to 95%, which is the efficacy of the vaccine for the wild type pathogen. The escape depth decreases faster towards 0 for a high efficacy of the vaccine towards the escape mutants than for low values, but independently of the exact value the escape depth will decrease to 0 for increasing number of vaccination types in the system.

Secondly, we evaluated the effect of the vaccine coverage in the model (see Figure 5**B**). The escape depth decreases fastest with the number of vaccination types when the vaccination coverage is high.

**Figure 5:**
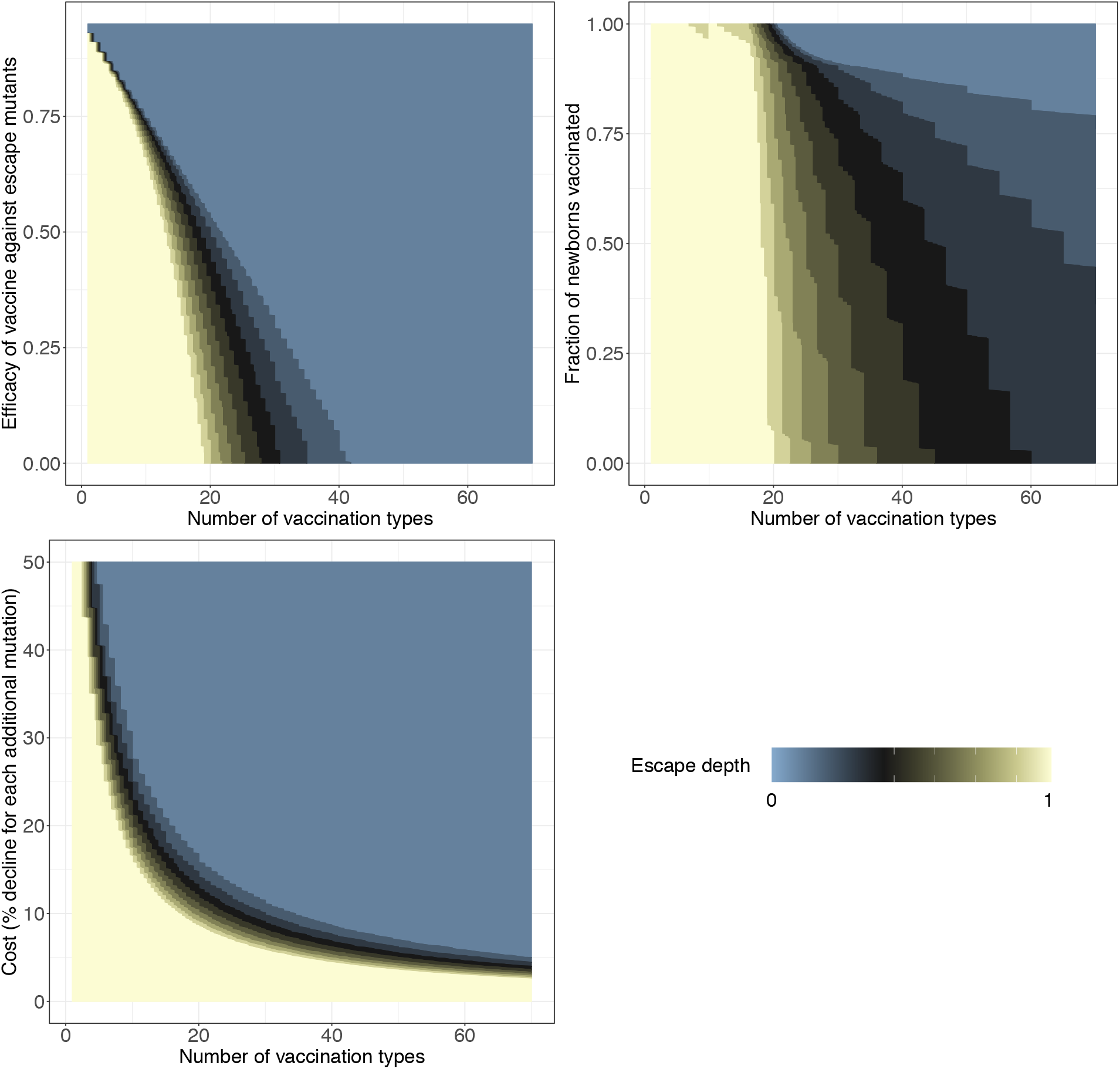
The effect of (A) the efficacy of the vaccine against escape mutants, (B) the fraction of newborns vaccinated, and (C) the mutation cost on the escape depth for an increasing number of vaccination types.

Thirdly, the cost of an escape mutation is varied, i.e. the decline in transmission rate per additional mutation. The escape depth decreases fastest when the cost of mutation is high. For low cost values (smaller than 2.5%), the escape depth does not decrease for the range of vaccination types we have evaluated.

Our model and analysis thus provide a rational and quantitative basis for evaluating the evolutionary sustainability of a vaccine, which can be predicted from its efficacy, coverage, and the fitness cost of escape.

### Within host multiplicity compared to between-host diversity of immune responses against epitopes

So far, we have considered vaccine-induced immune responses that only differ between individuals. However, it is known that a single host can respond against multiple antigens and epitopes of a pathogen [11], [12]. In a next step, we therefore consider vaccines that induce the same immune response in different individuals, but multiple responses within them. We assume that a mutant can only escape such an immune response fully if it has a mutation in each epitope, against which the vaccination-type responds, and partial escape mutants are as susceptible to the immune response as the wild-type while still carrying a transmission cost. Figure 6**A** shows how the basic reproduction number changes for an increasing number of within-host immune responses. We find that the basic reproduction number of partial escapes are unaffected by number of within-host epitopes. However, the basic reproduction number of the full escape mutant decreases when the number of within-host epitopes increases. This is due to the increased cost of the full escape mutant which is larger when the vaccine elicits more immune responses within a host.

**Figure 6:**
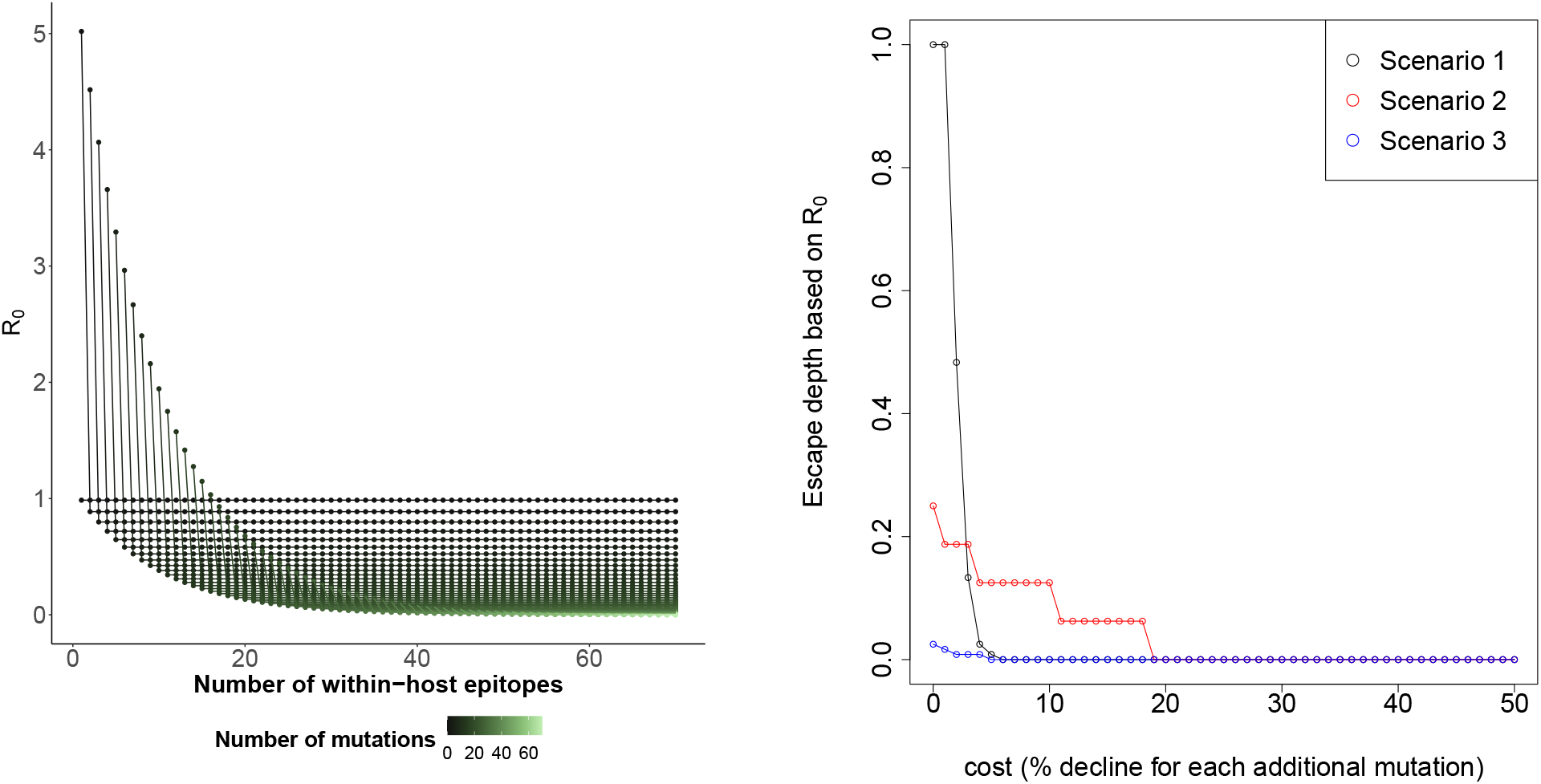
(A) The basic reproduction number for an increasing number of within-host epitopes to which the immune system responds upon vaccination in a model where there is no variation in responses between individuals. (B) The escape depth against the cost of an escape mutation for three different scenarios: one immune response against a single epitope per individual and 120 unique responses against epitopes in total (black); 14 different immune responses against epitopes per individual and 16 in the population (red); 119 immune responses against epitopes per individual and 120 immune responses in the population (blue).

Additionally, we tested the relative contribution of the within-host multiplicity of immune responses and the between-host diversity in preventing the spread of escape mutants. To this end, we construct three scenarios that cover the conceivable range of possibilities from extremely low to very high within-host multiplicity, while keeping the between-host diversity constant at 120 vaccination types. We then compare the escape depth in these three scenarios 6**B**).

The first scenario allows for one immune response against a single epitope per individual and 120 unique responses against epitopes in total. The second scenario has 14 different immune responses against epitopes per individual and 16 in the population, again resulting in 120 vaccination types. The third scenario has 119 immune responses against epitopes per individual and 120 immune responses in the population.

For low costs of escape, the vaccine with low within-host multiplicity, as described in the first scenario, has the highest, while the vaccine with high within-host multiplicity displays the lowest escape depth. Thus vaccines with low within-host multiplicity are predicted to be evolutionary least sustainable. This is due to the fact that for such vaccines a single mutation can already confer escape. Vaccines with intermediate within-host multiplicity show intermediate sustainability. When the mutation cost is high, the patterns changes: both, the vaccine with low and high within-host multiplicity are predicted to be most evolutionary sustainable. Thus, the escape depth, and hence the predicted evolutionary sustainability, is affected by a complex interaction between the cost of escape and the within-host multiplicity.

## Discussion

Consistent with previous studies, our model shows that the potential of an escape mutant to spread within a vaccinated population decreases with an increase in the diversity in immune responses upon vaccination, i.e. the number of vaccination-types in the population. We further show that, when the diversity in immune responses is large enough, the spread of escape mutants is predicted to be even completely preventable. This critical diversity depends on the cost of an escape mutation, the vaccination coverage and the efficacy of the vaccine against escape mutants. To summarize the complex escape dynamics in our model with multiple immune reponses and pathogen strains, we propose the new concept of the escape depth.

Additionally, our analyses showed that, for the prevention of vaccine escape, the between-host diversity of epitopes, to which the immune system responds, is more important than the multiplicity of these epitopes within hosts. However, one important effect of responding against multiple epitopes within a host is that it makes it harder for an escape mutant to emerge de novo, as it needs multiple mutations to escape the immune response of a single individual. This effect is not included in our model, because we assume that all escape mutants are already present in the population at low levels. Therefore, the advantage of having multiple immune responses with an individual is likely underestimated in our analysis.

Throughout the study, we have used the term *evolution proof* cautiously. Even for model scenarios where the predicted escape depth is zero, vaccine escape mutants could emerge and spread in reality. The reason for this is that escape mutants could pick up compensatory mutations that relieve the cost of the escape mutations, while we exclude such compensatory mutations in our simulations. Thus the cost of escape, which is central in our analysis of vaccine sustainability, could evolve to zero in the long run.

Our predictions agree with experimental findings involving the prokaryotic CRISPR-Cas system. Within this system, bacteria can acquire immunity against a pathogen, such as a bacteriophage, by inserting a spacer derived from the DNA of the pathogen into the CRISPR locus [23], [24]. Even for the same pathogen, this spacer can differ between bacterial cells [25]. Van Houte et al. (2016) showed that the probability that a pathogen goes extinct when invading a bacterial population increases with the spacer diversity [26]. Thus, our main finding, that diversity in immune responses decreases the probability that escape mutants spread, is supported in phage-bacteria systems.

In their inspiring perspective Kennedy and Read [1] list several examples of vaccine escape in systems with low immune diversity that are consistent with the epidemiological dynamics predicted by our model. First, the vaccine against Hepatitus B Virus (HBV) elicits immune responses against one epitope and a single mutation confers vaccination escape [27], [28]. As a result, vaccine escape mutants evolved shortly after the introduction of the vaccine [29]. Second, escape mutants of *Yersinia Ruckeri*, that causes Enteric redmouth disease (ERD) in fish, emerged after the introduction of a first monovalent vaccine [30]–[32]. This escape can be caused by four different single mutations which all effect the same pathway [1], [31]. On the other hand, one of the best-known examples of vaccination escape, Marek’s Disease Virus (MDV), which causes a viral infection in chicken [33], is not well described by our model. This is because the MDV strain used as a live vaccine, rather than inducing protective immune responses, works by occupying the target cell niche of the pathogenic MDV strain [34]. As a consequence, escaping from the MDV vaccine can be accomplished, not by avoiding the detection by immune responses, but by increasing the competitive ability against the vaccine strain [35].

In the current SARS-CoV-2 pandemic, there is concern that a vaccine resistant mutant already exists [36] or will emerge [37]. Our model uses realistic assumptions for a pandemic situation, as, due to the high case numbers world wide, we can assume that many possible mutants are around. The natural immune response against SARS-CoV-2 is diverse [38], therefore, immune responses upon vaccination might be diverse as well. Determining the diversity of responses following vaccination against COVID-19 will thus allow us to obtain a clearer picture about the sustainability of COVID-19 vaccines.

The question of why drug-resistant pathogens are so widely observed, while vaccination-escape mutants are not, has been addressed with mathematical models previously [1], [22]. With the present paper, we are contributing to this growing body of work by not only investigating the impact of immune diversity on the potential to delay the spread of escape mutants, but to prevent their spread altogether. Most crucial for the potential spread of escape mutants are their fitness costs. Thus, determining the diversity of vaccine-induced immune responses within individuals and the population, as well as the costs of escape mutations could allow us to better predict the sustainability of vaccines in the face of pathogen evolution.

## Methods and Materials

### The model

We use a mathematical model to investigate under which conditions the diversity in immune responses against epitopes can prevent the spread of vaccine escape mutants in a vaccinated population. Our model is an extension of McLean’s (1995) model [22]. The model starts with a susceptible, non-vaccinated population (*S*). A wild-type infectious pathogen can infect the susceptible population with transmission rate *β*. Individuals infected with the wild-type are indicated with *I^w^*. After twenty years, a vaccination program starts and 80 percent of the newborns are vaccinated with a vaccine that has an efficacy of 95 percent for the wild-type pathogen. There is also an escape mutant present in the system, for which the vaccine is only 20 percent efficient. Individuals infected with either type of pathogen who recover are immune to both pathogen types for the rest of the simulation.

We expanded McLean’s model to include diversity in immune responses upon vaccination. As indicated in Figure 1, each individual can elicit an immune response against a variety epitopes and each unique combination of immune responses against epitopes forms a vaccination-type. Figure 7 represents a model where six vaccination-types are possible. We assume that all immune responses against epitopes are equally distributed over the population. Besides the wild-type pathogen, there are also various escape mutants around. Each escape mutant contains a mutation in one ore multiple of the epitopes the immune system can recognize. In Figure 7, compartment *I* represents individuals that are infected with the wild-type or one of the escape mutants. All possible escape mutants exist before vaccination starts. The death rate of vaccinated and unvaccinated individuals is equal, as well as the rate at which individuals leave any of the infected compartments. The advantage of escaping any of the immune responses against epitopes comes at a cost. Therefore, we let the transmission rate at which a mutant can spread decay with 10 percent for each mutation. The full system of ODE equations that describes this model can be found in the Supplementary material, as well as a table with a description and the value of all model parameters.

**Figure 7:**
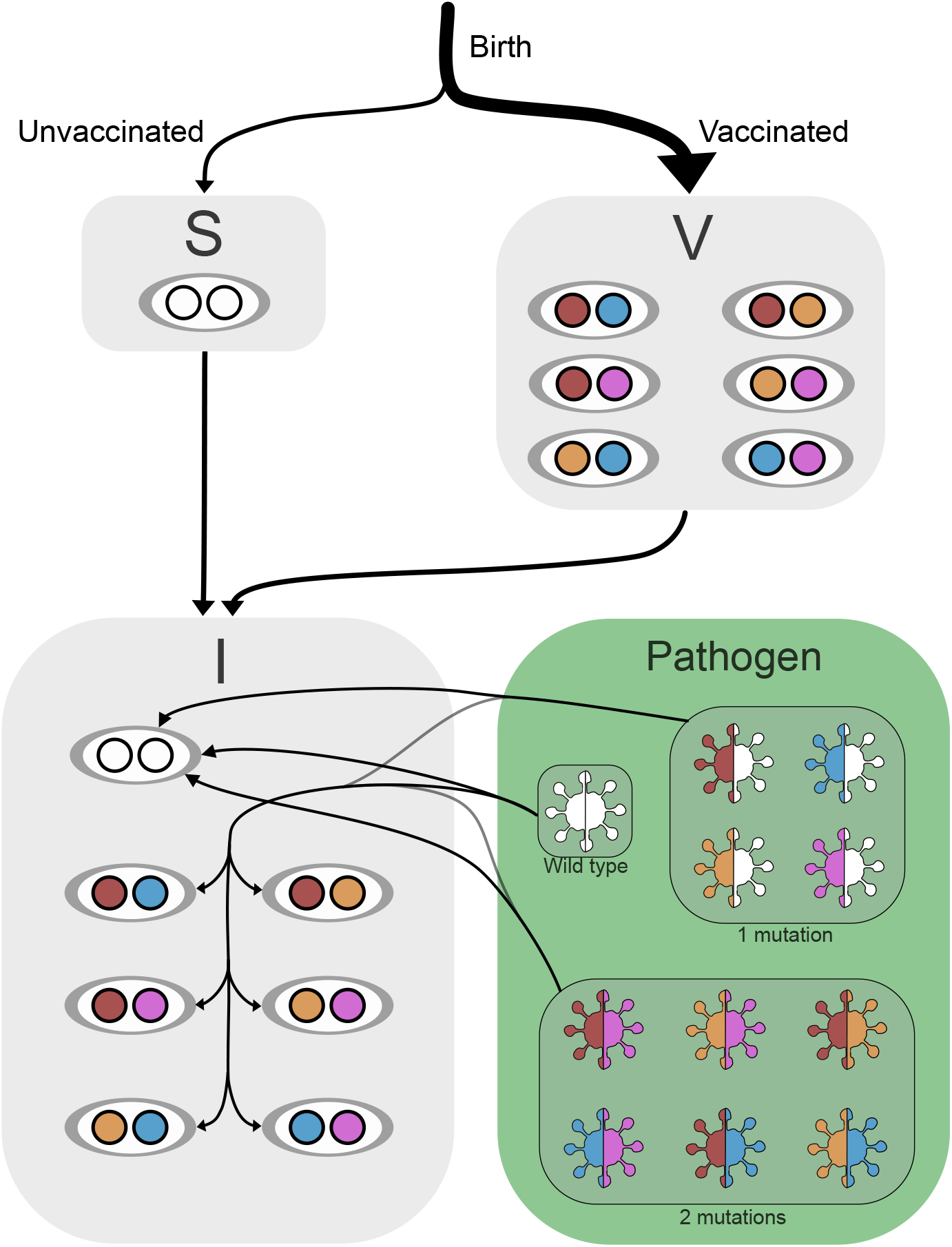
Schematic representation of the model when each individual can elicit an immune response against two different epitopes upon vaccination and there are four different epitopes in the system. *S* represents the compartment of unvaccinated susceptibles and *V* the vaccinated individuals that elicit one of the six vaccination-types. The compartment *Pathogen* shows the wild type virus and all possible escape mutants. *I* are individuals infected with either the wild-type pathogen or one of the escape mutants.

### Calculation of Reproduction Numbers

In our deterministic model, an escape mutant can only spread in the population, when it has a reproduction number larger than one. Therefore, we analyse the model to find analytical expressions as well as numerical values for the reproduction number of each escape mutant. We do this for both the reproduction number in a fully naive population, so without any other escape mutants around (the basic reproduction number *R*_0_), and in a situation where all other escape mutants that have less escape mutations where already able to spread (the endemic reproduction number *R_e_*). The general equation for both reproduction numbers is shown in Figure 3 and is a combination of the transmission rate of the pathogen type (*β*), the duration of the infectious period 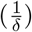 and the number of individuals that can be infected by this pathogen type. For the basic reproduction number, this number of individuals is calculated using the disease free equilibria for the number of vaccinated and non-vaccinated susceptibles: 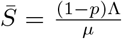 and 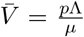For the endemic reproduction number of each pathogen type, we used the equilibria values of the number of non-vaccinated and vaccinated individuals in a system where all other pathogen types with less mutations have already spread. These cannot be calculated analytically and are therefore calculated numerically.

### Technical Implementation

The modelling was done in R (version 3.4.1) [39] using an automatized code that allows for an arbitrary number of vaccination-types and epitopes against which the immune system responds. All code is available on https://gitlab.ethz.ch/jbouman/Vaccine_escape_open. The deSolve package [40] is used to numerically solve the system of differential equations. Additionally the function *as.set* from the *sets* package is used to create a set with all possible escape mutants [41]. Endemic equilibria are found by solving the ODE system for its roots, using the *multiroot-function* from *rootSolve* [42], [43].

## Acknowledgement

The authors thank Claudia Igler and Michael Manhart for helpful comments.

## Supplementary Material

### Full Model Description

The model depicted in Figure 7 is described by the equations below for one vaccination-type per individual. A description of all parameters and their value used for the simulations is shown in Table.

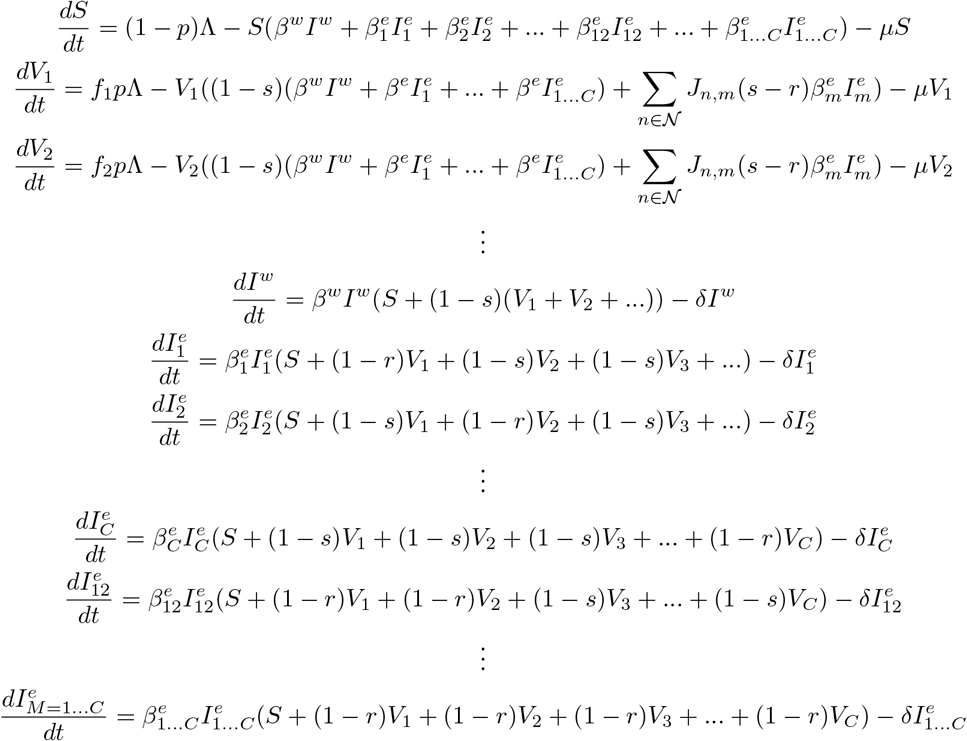

The set of equations that describes the complete model. The meaning of the different parameters is described in table. 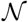 is the set that contains all vaccination-types, *J_n,m_* equals one if mutant *m* can escape vaccination-type 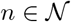 and otherwise zero.

---

**Table S1:** Description of all model parameters and their values.

| Parameter | Meaning | Value |
| --- | --- | --- |
| $S$ | Number of non-vaccinated susceptibles | Initial value: 9000 |
| $V_i$ | Number of vaccinated susceptibles that elicit vaccination-type $i$ | Initial value: 0 |
| $I^w$ | Number of individuals infected with the wild-type pathogen | Initial value: 10 |
| $I_i^e$ | Number of individuals infected with an escape mutant of type $i$ | Initial value: 1 for each type $i$ |
| $p$ | Fraction of newborn susceptibles that will be vaccinated starting after three years of the simulation | 0.9 |
| $\Lambda$ | Birth rate | 2000 per year |
| $\beta^w$ | Transmission rate for the wild-type pathogen | 0.0017 |
| $\beta_i^e$ | Transmission rate for the escape mutant indicated with $i$ | Dependent on scenario |
| $f_i$ | Fraction of vaccinated individuals that elicits vaccination-type $i$ | $\frac{1}{\text{number of vaccination-types}}$ |
| $s$ | efficacy of vaccination for the wild-type pathogen | 0.95 |
| $r$ | efficacy of vaccination on pathogen that can escape this vaccination-type | 0.2 |
| $\mu$ | Death rate for vaccinated and non-vaccinated susceptible population | 0.02 per year |
| $\delta$ | Rate at which individuals leave any of the infected compartments (due to death or recovery) | 25 per year |

## Supplementary Result Figures

**Figure S1:**
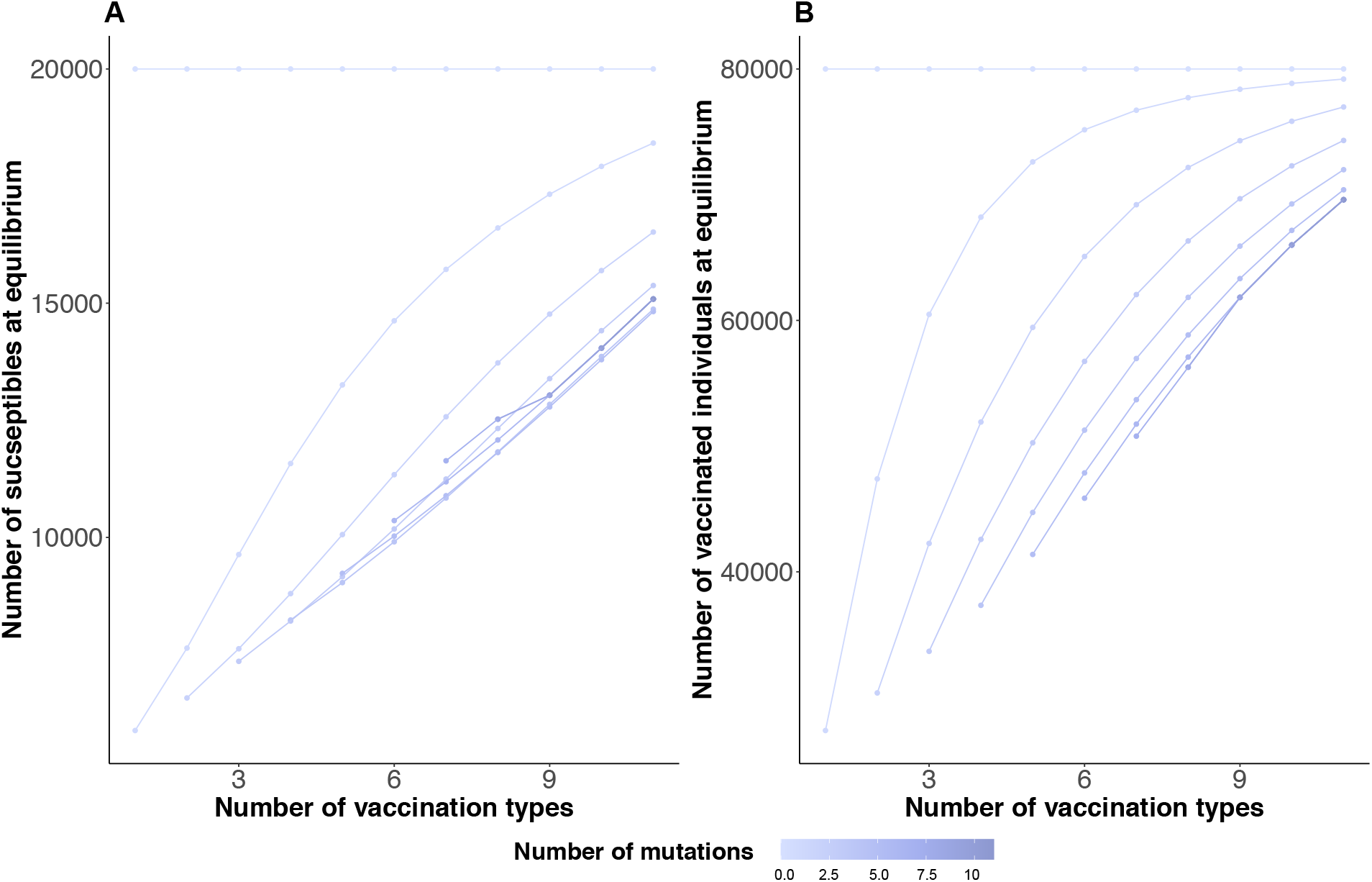
(A) The number of susceptible individuals (non-vaccinated) at the equilibrium of the system with varying number of vaccination types and for a selection of mutants available in the system. (B) The number of susceptible vaccinated individuals at equilibrium.

## References

[1] D. A. Kennedy and A. F. Read, “Why does drug resistance readily evolve but vaccine resistance does not?” Proceedings of the Royal Society B: Biological Sciences, vol. 284, no. 1851, p. 20 162 562, 2017.

[2] P. A. zur Wiesch, R. Kouyos, J. Engelstädter, R. R. Regoes, and S. Bonhoeffer, “Population biological principles of drug-resistance evolution in infectious diseases,” The Lancet infectious diseases, vol. 11, no. 3, pp. 236–247, 2011.

[3] J. Davies and D. Davies, “Origins and evolution of antibiotic resistance,” Microbiol. Mol. Biol. Rev., vol. 74, no. 3, pp. 417–433, 2010.

[4] M. Barber and J. Whitehead, “Bacteriophage types in penicillin-resistant staphylococcal infection,” British medical journal, vol. 2, no. 4627, p. 565, 1949.

[5] M. F. Flajnik and M. Kasahara, “Origin and evolution of the adaptive immune system: Genetic events and selective pressures,” Nature Reviews Genetics, vol. 11, no. 1, pp. 47–59, 2010.

[6] J. Travis, On the origin of the immune system, 2009.

[7] B. R. Bloom and P.-H. Lambert, The vaccine book. Academic Press, 2002.

[8] M. B. Doud, J. M. Lee, and J. D. Bloom, “How single mutations affect viral escape from broad and narrow antibodies to h1 influenza hemagglutinin,” Nature communications, vol. 9, no. 1, pp. 1–12, 2018.

[9] T. T. Immonen, C. Camus, C. Reid, C. M. Fennessey, G. Q. Del Prete, M. P. Davenport, J. D. Lifson, and B. F. Keele, “Genetically barcoded siv reveals the emergence of escape mutations in multiple viral lineages during immune escape,” Proceedings of the National Academy of Sciences, vol. 117, no. 1, pp. 494–502, 2020.

[10] J. D. Galson, A. J. Pollard, J. Trück, and D. F. Kelly, “Studying the antibody repertoire after vaccination: Practical applications,” Trends in immunology, vol. 35, no. 7, pp. 319–331, 2014.

[11] J. J. Lavinder, Y. Wine, C. Giesecke, G. C. Ippolito, A. P. Horton, O. I. Lungu, K. H. Hoi, B. J. DeKosky, E. M. Murrin, M. M. Wirth, et al., “Identification and characterization of the constituent human serum antibodies elicited by vaccination,” Proceedings of the National Academy of Sciences, vol. 111, no. 6, pp. 2259–2264, 2014.

[12] T. R. Poulsen, P.-J. Meijer, A. Jensen, L. S. Nielsen, and P. S. Andersen, “Kinetic, affinity, and diversity limits of human polyclonal antibody responses against tetanus toxoid,” The Journal of Immunology, vol. 179, no. 6, pp. 3841–3850, 2007.

[13] K. Smith, J. J. Muther, A. L. Duke, E. McKee, N.-Y. Zheng, P. C. Wilson, and J. A. James, “Fully human monoclonal antibodies from antibody secreting cells after vaccination with pneumovax® 23 are serotype specific and facilitate opsonophagocytosis,” Immunobiology, vol. 218, no. 5, pp. 745–754, 2013.

[14] J. J. Goronzy and C. M. Weyand, “T cell development and receptor diversity during aging,” Current opinion in immunology, vol. 17, no. 5, pp. 468–475, 2005.

[15] C. Giefing-Kröll, P. Berger, G. Lepperdinger, and B. Grubeck-Loebenstein, “How sex and age affect immune responses, susceptibility to infections, and response to vaccination,” Aging cell, vol. 14, no. 3, pp. 309–321, 2015.

[16] S. L. Klein, A. Jedlicka, and A. Pekosz, “The xs and y of immune responses to viral vaccines,” The Lancet infectious diseases, vol. 10, no. 5, pp. 338–349, 2010.

[17] G. A. Poland, R. M. Jacobson, S. A. Colbourne, A. M. Thampy, J. J. Lipsky, P. C. Wollan, P. Roberts, and S. J. Jacobsen, “Measles antibody seroprevalence rates among immunized inuit, innu and caucasian subjects,” Vaccine, vol. 17, no. 11-12, pp. 1525–1531, 1999.

[18] B. Posteraro, R. Pastorino, P. Di Giannantonio, C. Ianuale, R. Amore, W. Ricciardi, and S. Boccia, “The link between genetic variation and variability in vaccine responses: Systematic review and meta-analyses,” Vaccine, vol. 32, no. 15, pp. 1661–1669, 2014.

[19] R. Kurupati, A. Kossenkov, L. Haut, S. Kannan, Z. Xiang, Y. Li, S. Doyle, Q. Liu, K. Schmader, L. Showe, et al., “Race-related differences in antibody responses to the inactivated influenza vaccine are linked to distinct pre-vaccination gene expression profiles in blood,” Oncotarget, vol. 7, no. 39, p. 62 898, 2016.

[20] J. Glanville, T. C. Kuo, H.-C. von Büdingen, L. Guey, J. Berka, P. D. Sundar, G. Huerta, G. R. Mehta, J. R. Oksenberg, S. L. Hauser, et al., “Naive antibody gene-segment frequencies are heritable and unaltered by chronic lymphocyte ablation,” Proceedings of the National Academy of Sciences, vol. 108, no. 50, pp. 20 066–20 071, 2011.

[21] C. Wang, Y. Liu, M. M. Cavanagh, S. Le Saux, Q. Qi, K. M. Roskin, T. J. Looney, J.-Y. Lee, V. Dixit, C. L. Dekker, et al., “B-cell repertoire responses to varicella-zoster vaccination in human identical twins,” Proceedings of the National Academy of Sciences, vol. 112, no. 2, pp. 500–505, 2015.

[22] A. R. McLean, “Vaccination, evolution and changes in the efficacy of vaccines: A theoretical framework,” Proceedings of the Royal Society of London. Series B: Biological Sciences, vol. 261, no. 1362, pp. 389–393, 1995.

[23] R. Barrangou, C. Fremaux, H. Deveau, M. Richards, P. Boyaval, S. Moineau, D. A. Romero, and P. Horvath, “Crispr provides acquired resistance against viruses in prokaryotes,” Science, vol. 315, no. 5819, pp. 1709–1712, 2007.

[24] J. Van Der Oost, E. R. Westra, R. N. Jackson, and B. Wiedenheft, “Unravelling the structural and mechanistic basis of crispr–cas systems,” Nature Reviews Microbiology, vol. 12, no. 7, pp. 479–492, 2014.

[25] D. Paez-Espino, W. Morovic, C. L. Sun, B. C. Thomas, K.-i. Ueda, B. Stahl, R. Barrangou, and J. F. Banfield, “Strong bias in the bacterial crispr elements that confer immunity to phage,” Nature communications, vol. 4, no. 1, pp. 1–7, 2013.

[26] S. van Houte, A. K. Ekroth, J. M. Broniewski, H. Chabas, B. Ashby, J. Bondy-Denomy, S. Gandon, M. Boots, S. Paterson, A. Buckling, et al., “The diversity-generating benefits of a prokaryotic adaptive immune system,” Nature, vol. 532, no. 7599, pp. 385–388, 2016.

[27] L. Romanò, S. Paladini, C. Galli, G. Raimondo, T. Pollicino, and A. R. Zanetti, “Hepatitis b vaccination: Are escape mutant viruses a matter of concern?” Human vaccines & immunotherapeutics, vol. 11, no. 1, pp. 53–57, 2015.

[28] J. Sheldon and V. Soriano, “Hepatitis b virus escape mutants induced by antiviral therapy,” Journal of antimicrobial chemotherapy, vol. 61, no. 4, pp. 766–768, 2008.

[29] R Zanetti, E Tanzi, G Manzillo, G Maio, C Sbreglia, N Caporaso, H. Thomas, A. Zuckerman, et al., “Hepatitis b variant in europe,” 1988.

[30] R. Busch, “Protective vaccines for mass immunization of trout,” Salmonid, vol. 1, no. 6, pp. 10–14, 1978.

[31] D. A. Austin, P. Robertson, and B. Austin, “Recovery of a new biogroup of yersinia ruckeri from diseased rainbow trout (oncorhynchus mykiss, walbaum),” Systematic and Applied Microbiology, vol. 26, no. 1, pp. 127–131, 2003.

[32] T. J. Welch, D. W. Verner-Jeffreys, I. Dalsgaard, T. Wiklund, J. P. Evenhuis, J. A. G. Cabrera, J. M. Hinshaw, J. D. Drennan, and S. E. LaPatra, “Independent emergence of yersinia ruckeri biotype 2 in the united states and europe,” Appl. Environ. Microbiol., vol. 77, no. 10, pp. 3493–3499, 2011.

[33] H. K. Adldinger and B. W. Calnek, “Pathogenesis of marek’s disease: Early distribution of virus and viral antigens in infected chickens,” Journal of the National Cancer Institute, vol. 50, no. 5, pp. 1287–1298, 1973.

[34] L. Yakovleva and N. Mazurenko, “Notes on the mechanism of postvaccination immunity in marek’s disease.,” Neoplasma, vol. 24, no. 4, pp. 387–394, 1977.

[35] R. Witter, “Avian tumor viruses: Persistent and evolving pathogens.,” Acta Veterinaria Hungarica, vol. 45, no. 3, pp. 251–266, 1997.

[36] J. Wise, “Covid-19: The e484k mutation and the risks it poses,” 2021.

[37] Y. H. G. Sarah Cobey Daniel B. Larremore and M. Lipsitch, “Concerns about sars-cov-2 evolution should not hold back efforts to expand vaccination,” 2021.

[38] Y. Peng, A. J. Mentzer, G. Liu, X. Yao, Z. Yin, D. Dong, W. Dejnirattisai, T. Rostron, P. Supasa, C. Liu, et al., “Broad and strong memory cd4+ and cd8+ t cells induced by sars-cov-2 in uk convalescent individuals following covid-19,” Nature immunology, vol. 21, no. 11, pp. 1336–1345, 2020.

[39] R. C. Team, “R: A language and environment for statistical com-puting,” R Foundation for Statistical Computing, Vienna, Austria. URL https://www.R-project.org, 2017.

[40] K. E. Soetaert, T. Petzoldt, and R. W. Setzer, “Solving differential equations in r: Package desolve,” Journal of Statistical Software, vol. 33, 2010.

[41] K. Hornik and D. Meyer, “Generalized and customizable sets in r,” Journal of Statistical Software, vol. 31, no. 2, pp. 1–27, 2009.

[42] K Soetaert and P. Herman, “Ecolmod:” a practical guide to ecological modelling-using r as a simulation platform”,” R package version, vol. 1, 2009.

[43] K. Soetaert, “Rootsolve: Nonlinear root finding, equilibrium and steady-state analysis of ordinary differential equations,” R package version, vol. 1, no. 6, 2009.

